# Dynamic optimization reveals alveolar epithelial cells as key mediators of host defense in invasive aspergillosis

**DOI:** 10.1101/2021.05.12.443764

**Authors:** Jan Ewald, Flora Rivieccio, Lukáš Radosa, Stefan Schuster, Axel A. Brakhage, Christoph Kaleta

## Abstract

*Aspergillus fumigatus* is an important human fungal pathogen and its conidia are constantly inhaled by humans. In immunocompromised individuals, conidia can grow out as hyphae that damage lung epithelium. The resulting invasive aspergillosis is associated with devastating mortality rates. Since infection is a race between the innate immune system and the outgrowth of *A. fumigatus* conidia, we use dynamic optimization to obtain insight into the recruitment and depletion of alveolar macrophages and neutrophils. Using this model, we obtain key insights into major determinants of infection outcome on host and pathogen side. On the pathogen side, we predict *in silico* and confirm *in vitro* that germination speed is a key virulence trait of fungal pathogens due to the vulnerability of conidia against host defense. On the host side, we find that epithelial cells play a so far underappreciated role in fungal clearance and are potent mediators of cytokine release which we confirm *ex vivo*. Further, our model affirms the importance of neutrophils in invasive aspergillosis and underlines that the role of macrophages remains elusive. We expect that our model will contribute to improvement of treatment protocols by focusing on the critical components of immune response to fungi but also fungal virulence.

## 1. Introduction

Since we constantly inhale microorganisms, the human lung is an entry point for opportunistic pathogenic microorganisms, like the mould *Aspergillus fumigatus* [1, 2]. Besides being a saphrophyte involved in the decay of organic matter in soil, *A. fumigatus* possesses virulence characteristics such as small spores (conidia), a fast growth at body temperature and the production of specific proteins, carbohydrates and secondary metabolites allowing for immune evasion [3–7]. These traits enable *A. fumigatus* to reach the lung alveoli and cause an invasive aspergillosis by filamentous growth (hyphae) into the tissue and dissemination into the host [5, 8]. In the immunocompetent host this is prevented by the fast and efficient clearance of conidia by the innate immune system within a few hours [7, 9]. However, once *A. fumigatus* grows invasively into human tissues facilitated by a suppressed immune system, mortality is very high (30-95%) due to non-efficient diagnostics and limited treatment options [10–13].

Along with the advances in medical care the number of patients with defects and suppression of their immune system is expected to grow. Major causes for this trend are the increasing number of cancer patients receiving chemotherapy [14, 15], organ transplant recipients [16] or patients with acquired immune deficiency syndrome (AIDS) [17]. Most recently, a high number of COVID-19 patients in intensive care units with extensive ventilation has been accompanied by secondary fungal infections due to *Aspergillus spp*. [13, 18].

The race between fungal growth and host immune response is complex and involves many cells like alveolar epithelial cells (AEC), alveolar macrophages (AM) and recruited neutrophils [7, 9]. To better understand this spatial and dynamic process computational modeling has proven to be of value [19–24]. These models are based on like agent-based modelling or differential equation systems and contribute to a better understanding of the immune response. In particular, the meaning of spatial-dynamics of AM clearing conidia [20, 22, 23, 25] as well as the influence of the initial fungal burden on clearance and persistence of *A. fumigatus* were studied [19]. A major achievement and advantage of *in silico* models is the integration of existing biological knowledge and the generation of new hypotheses by disclosing knowledge gaps.

Despite extensive modeling and experimental investigations, the relative contribution of individual host immune cell types in invasive aspergillosis remains elusive [8]. For example, AM were identified as phagocytes of conidia [26] and release cytokines upon fungal infection [27]. Yet, AM depletion in mice at early phases of infection showed no effect on mortality while neutrophil depletion was accompanied with low survival rates [28]. Current models and their analysis are based on immune cells and virtually neglect the contribution of AEC to cytokine release and fungal clearance. Experimental data obtained *in vitro*, however, suggest that AEC can act as potent phagocytes [29] and release a significant amount of cytokines during their interaction with fungal conidia [30]. It has been postulated that AEC represent a ’neglected portal entry of *Aspergillus*’ [7]. Experimental studies exploring their meaning and functions for invasive aspergillosis are difficult to perform due to the multicellular interactions required to evaluate the role of AEC. Therefore, a focus of our presented model here, is to dissect the contributions of AEC and innate immune cells for defense against *A. fumigatus*.

To elucidate the dynamic process of invasive aspergillosis during the first 24h, we propose a model using dynamic optimization as a mathematical approach. The mathematical concept of a dynamic system described by differential equations and regulated by control variables matches the dynamics of the innate immune response with recruitment and depletion of neutrophils and AM as monocytes. Due to this advantage, dynamic optimization has also been used to model other host-pathogen interactions and immune responses [24, 31, 32] and makes use of the fact that these energy demanding processes are highly optimized during evolution [33]. Our presented model, in addition, not only elucidates the complex recruitment dynamics of immune cells, but we also study the role of AEC during early stages of fungal infection. To proof key variables of the model and thus its validity, major findings like fungal germination and cytokine release were experimentally evaluated.

## 2. Results

### 2.1. Model overview

The aim and scope of our modeling are a better understanding of the decisive parameters and interactions contributing to infection by *A. fumigatus* in the first 24h. To this end, we model the different growth states of *A. fumigatus* and their interaction with the innate immune response in the lung alveoli during the first hours of infection (see Fig. 1). In addition, we explicitly model AEC as interactive cells and consider a single dosage scenario of fungal conidia exposure. The latter modeling decision enables comparison of results and parameters to experimental animal models, which mainly use single dosage regimes [34, 35].

**Figure 1:**
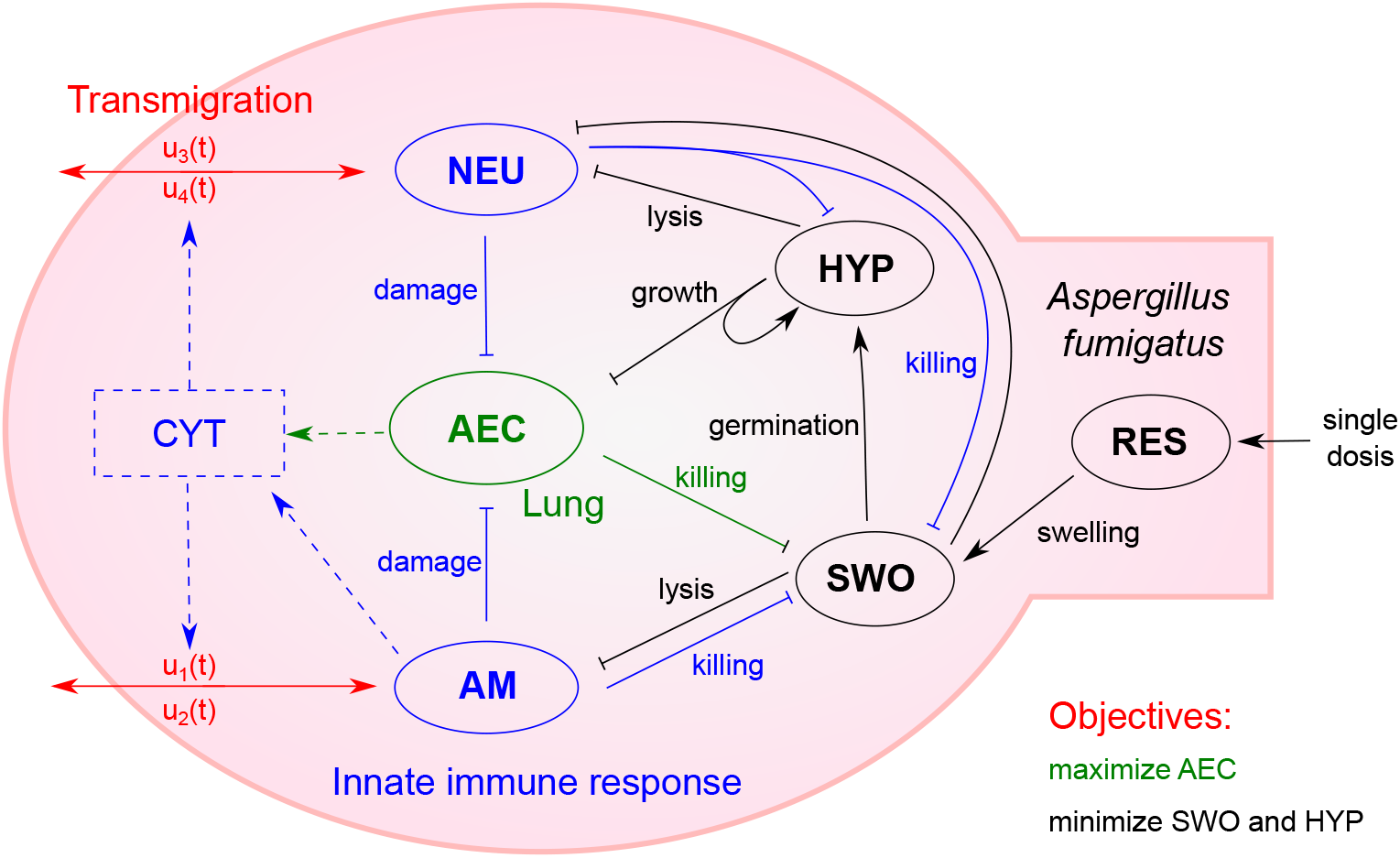
Model of invasive aspergillosis and the innate immune response as dynamic optimization problem. The different fungal growth states (black) including resting conidia (RES), swollen conidia (SWO) and hyphae (HYP) interact with alveolar epithelial cells (AEC, green) as well as with neutrophils (NEU, blue) or alveolar macrophages (AM, blue). Immune cell population is optimized via transmigration (red), which is linked to the cytokine level (CYT, blue), and optimal recruitment and depletion attain the trade-off between pathogen minimization and tissue integrity. Arrow heads indicate a positive interaction and bars show negative interactions.

Our model based on ordinary differential equations (ODE) considers time-dependent transition kinetics to describe the process of swelling and germination of conidia (detailed description in Subsection 4.1). After germination, growth of *A. fumigatus* as hyphae depends on the presence of AEC as resource. Importantly, we accurately model interaction of each cell type with all host cell types. Resting conidia are not recognized or killed due to their coating and swollen conidia are phagocytosed and killed by AM, neutrophils and AEC. Since AM and AEC are unable to phagocytose and kill larger hyphae at reasonable rates [36, 37], we only model killing of hyphae by neutrophils.

Host cell dynamics are characterized by damage, host or pathogen mediated, and transmigration of immune cells. In our dynamic optimization model, AEC face lysis by hyphae and damage by activated immune cells. While AM and neutrophils like AEC show cell death upon interaction with fungal cells, their transmigration upon infection is modeled by recruitment and depletion. The maximal rate of recruitment and depletion is linked to the presence of pro-inflammatory cytokines and is optimized in our dynamic optimization approach via the control variables *u*_1–4_ (see Figure 1 and *cf*. Material and Methods 4.1). In our model we capture the release of cytokines by AM as well as AEC to reveal their respective contribution during infection.

As host objectives we define two main goals, which are optimized during infection. Firstly, active fungal cells (swollen conidia and hyphae) should be minimized at all time points to avert systemic infection. Secondly, unnecessary tissue damage *e.g.* by hyperinflammation and collateral damage mediated by immune cells must be avoided. The consideration of only one of the objectives leads to undesired dynamics like continued hyphae growth or extensive tissue damage, as shown in Appendix C. Hence, we performed optimization with an equal weighting of both objectives.

### 2.2. Parameters and time course of early immune response

For our dynamic optimization model of the innate immune response during early invasive aspergillosis, reference parameters were estimated based on an extensive study of existing data and literature (see Appendix A). The model focuses on the events in the alveoli during the first 24h and we normalize all cell populations per alveolus. Experimental studies and data reveal that the average number of AEC is 11.4 (type I and II combined) in a healthy mouse [38, 39] and on average in every third or fourth alveolus an AM, respectively a neutrophil, is resident [40]. In animal models of invasive aspergillosis typically initial conidia dosages of 10^5^ to 10^7^ are applied as single dose [7, 35]. Since there are around 2.3 · 10^6^ alveoli in a murine lung [39] and not all conidia reach an alveolus, one resting conidium per alveolus relates to a typical fungal burden during experiments.

To elucidate the general pattern of innate immune response during invasive aspergillosis, we simulated 500 randomized parameter sets, where each parameter follows a log-normal distribution with the estimated reference value as the mode (maximum of distribution). The calculated time courses of the dynamic optimization reveal that healthy mice are able to clear even high fungal burden without a complete destruction of the epithelial cell barrier (see Figure 2 and details of solving the dynamic optimization in Subsection 4.2). This is achieved by a rapid cytokine release by AM as well as AEC and recruitment of neutrophils after conidial swelling. After 10 to 15h, recruitment of neutrophils is stopped and the immune cells are depleted to avoid unnecessary tissue damage. In this regard, our model well reflects experimental observations [41] and depicts the trade-off between pathogen clearance and tissue damage by the innate immune response. In our model, interestingly, AM are not recruited in great quantities and are mostly depleted after germination of conidia. Based on our estimated parameters of the kinetic rates, this is mostly due to slower killing of conidia compared to neutrophils and the ability of AEC to release cytokines in great quantities.

**Figure 2:**
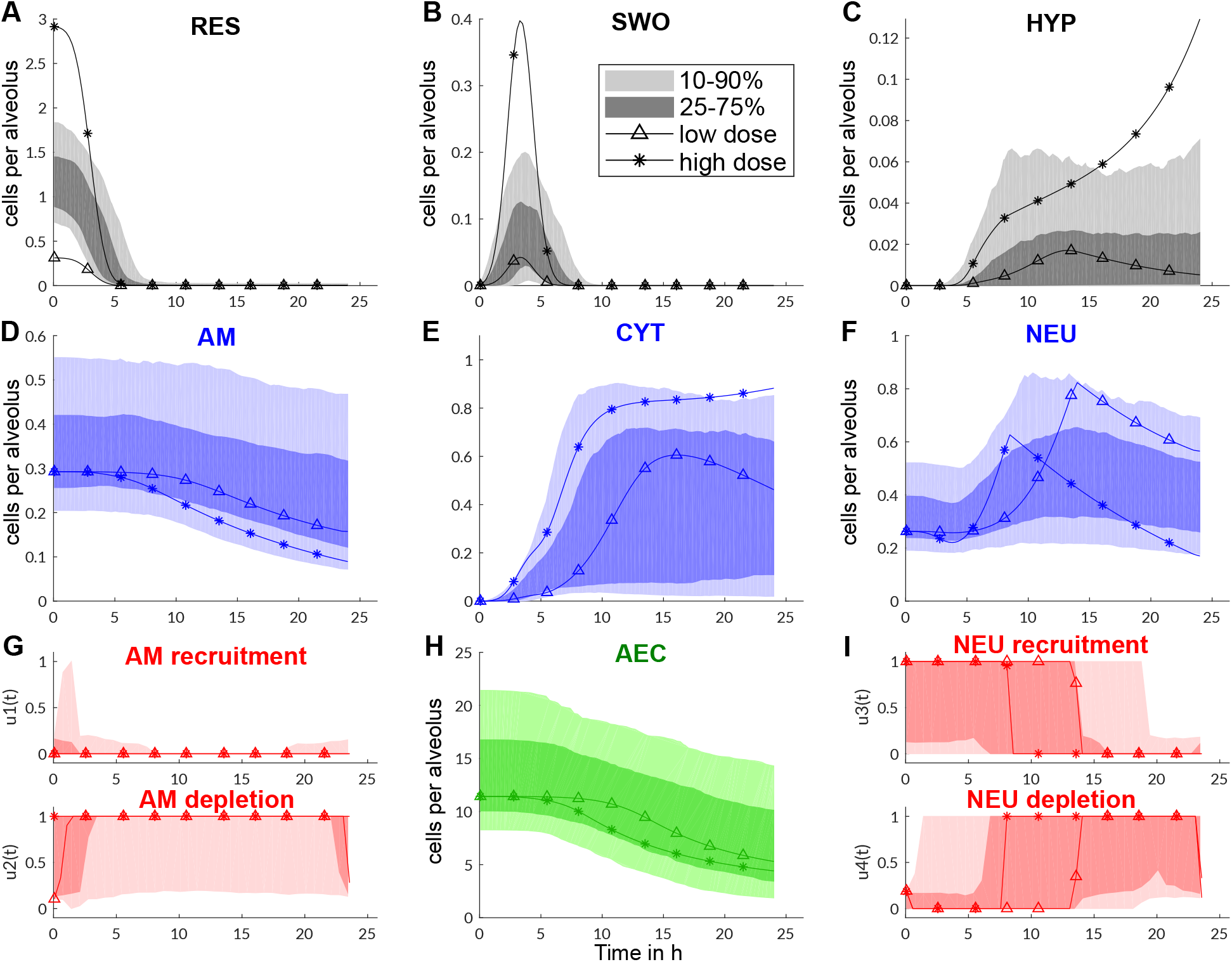
Dynamics of innate immune response of the murine host for varying parameters (shadings) and two dosage scenarios of conidia based on the reference parameter set (lines). **A-C** Dynamics of fungal cells per alveolus. **D, F** show the dynamics of the immune cells per alveolus which are influenced by the optimized recruitment and depletion rates in panels **G** and **I** (rates between 1, maximal, and 0, no recruitment/depletion). The recruitment and depletion potential is dependent on the cytokine level (**E**). Cytokines are produced by alveolar macrophages (**D**) and alveolar epithelial cells (**H**) in response to swollen conidia (**B**) or hyphae (**C**). The simulations of 500 parameter sets are depicted with shadings indicating the confidence intervals of time courses.

### 2.3. Parameter sensitivity reveals importance of fungal growth parameters

To better understand the roles of immune cells and AEC, we performed an analysis of parameter sensitivity and simulated several scenarios to study the effect of immunodeficiencies and conidial dosage.

We determined decisive parameters for the outcome of infection by calculating the contribution to variance in the objective function of each randomized parameter (see Material and Methods 4.2). Further, we simulated healthy mice and a lack of immune cells as well as the influence of low or high doses of conidia. Across all scenarios we see that the fungal parameters *s*_1_ (germination time of a swollen conidium) and *h*_1_ (hyphal growth rate) are most decisive for infection outcome since they explain more than 50% of the variance in the objective function (see Figure 3). This illustrates that a fast germination of swollen conidia is a strong virulence factor, because it is the most vulnerable growth state of the pathogen. The growth rate of hyphae is in addition crucial in the race between neutrophils (recruitment and hyphal killing) and *A. fumigatus*.

**Figure 3:**
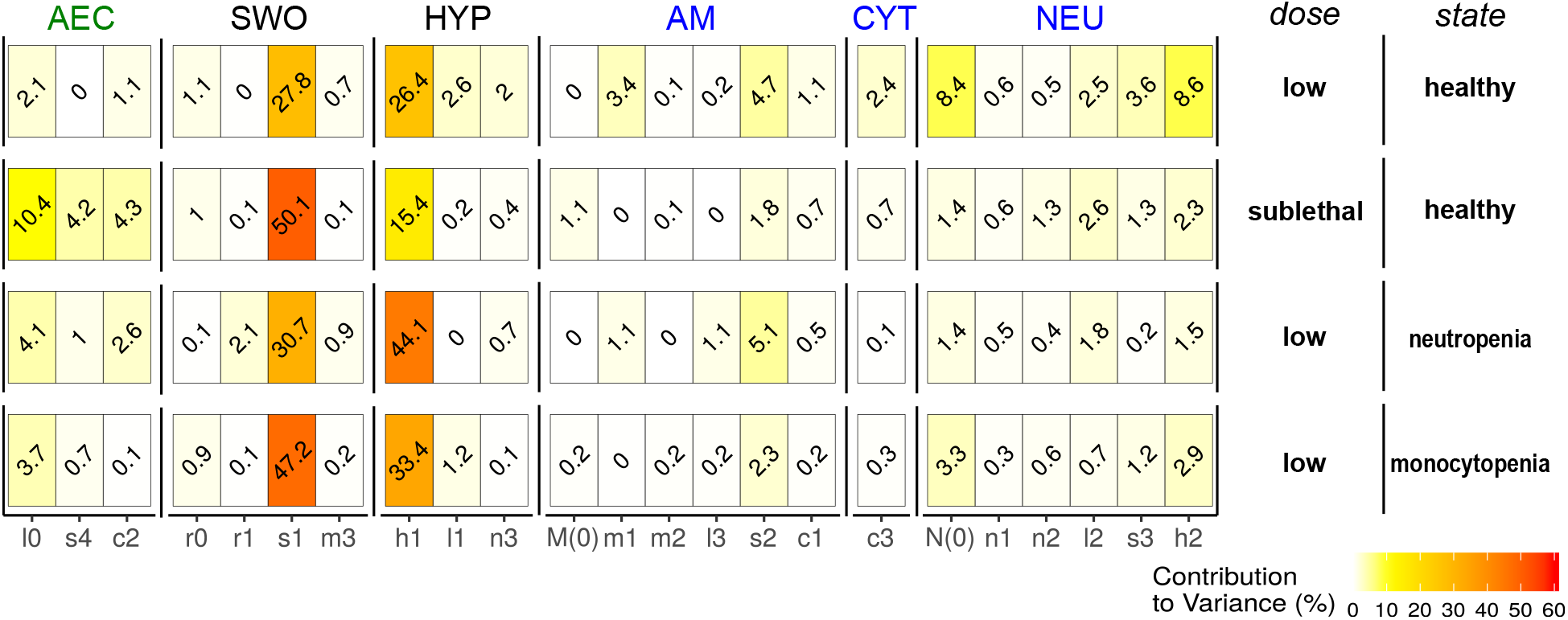
Influence of parameters on the outcome of infection depicted by the contribution to variance (colored from white, yellow to red from no to high influence). This relative contribution is based on a Spearman rank correlation of the parameter value and the objective value of the optimal solution (see Material and Methods 4.2). Parameters are grouped based on their relation to the cell types: alveolar epithelial cells (AEC), resting (RES) or swollen conidia (SWO), hyphae (HYP), alveolar macrophages (AM) and neutrophils (NEU). We list in addition the cytokine (CYT) property *c*_3_ (decay rate).

On the host side, we see that the importance of some parameters differs significantly when the initial fungal burden is altered. Interestingly, at lower dosages immune cell parameters like the number of resident neutrophils *N*(0) or rate of hyphal killing *h*_2_ are important (see Figure 3). However, at higher dosages parameters of AEC, such as conidia phagocytosis (*s*_4_) or cytokine release (*c*_2_), are more important for the infection outcome and contribute nearly 20% to the variance in the objective function. The parameter sensitivity and time course of infection indicate that at low dosages the control of hyphal growth is most important, while at high dosages AEC are crucial to lower the number of swollen conidia. These results are noteworthy since parameters related to functions of AM show even combined only a minor influence on the infection outcome (3.7% at high dosage and 9.5% at low dose). Typically, AM are extensively studied since they belong to the first line of defense. However, our parameter sensitivity results do not disclose a singular and distinctive role and we therefore performed additional analyses in the following sections.

Further insights can also be gained by studying the changes in parameter influence under scenarios of immunodeficiencies like the lack of monocytes (AM) or neutrophils simulated as reduced, 1%, immune cell populations and recruitment rates (see Figure 3). In both scenarios of immunodeficiencies, fungal virulence factors are even more important for infection outcome. But there are different tendencies as to which factor is more important. During monocytopenia the time span for germination, *s*_1_, is more important, suggesting that AM are mainly involved in the control of swollen conidia (see Figure 3). In contrast, during neutropenia the hyphal growth rate is more decisive for the infection outcome (see Figure 3) supporting the observation that neutrophils are crucial to prevent filamentous growth.

To support the findings established by modeling, we performed an experimental investigation of fungal virulence parameters by the comparison of different *Aspergillus* species. As revealed by the parameter sensitivity analysis, a key virulence factor is a fast germination to minimize the time period of the vulnerable swollen-conidia state. The common species *A. fumigatus, A. nidulans, A. niger* and *A. terreus* show differences in their germination kinetics at 37°*C*, where *A. nidulans* is fastest, *A. fumigatus* as well as *A. niger* are around 1-2h slower and *A. terreus* is by far the slowest with an average germination time of > 24*h* (see Figure 4A and B). Our model predicts here a non-linear but distinctive relationship between the germination time and epithelial damage after 24h (see Figure 4C). It suggests that virulence expressed as epithelial damage is strongly reduced if the germination time is longer than 10h.

**Figure 4:**
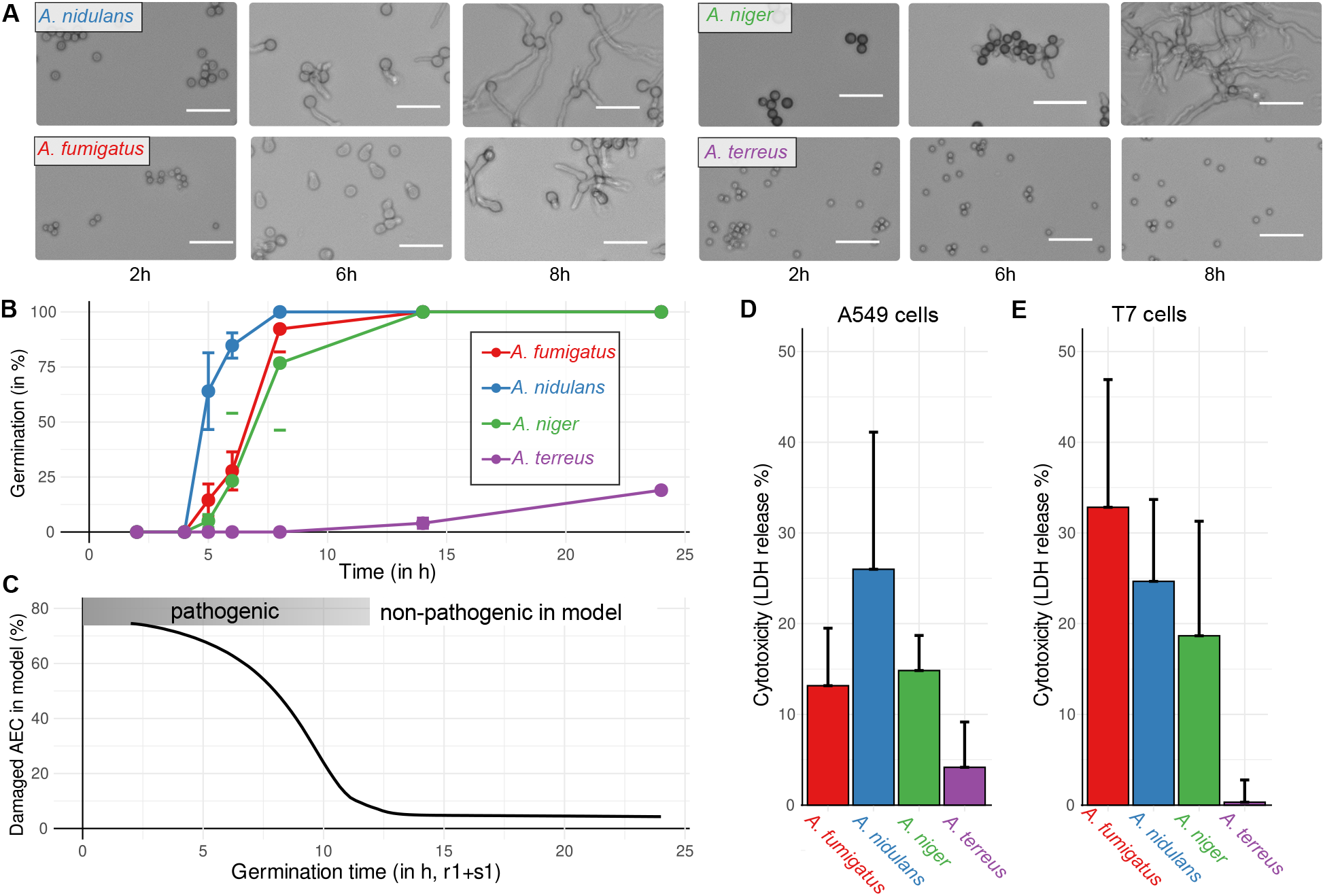
Germination kinetics of *Aspergillus spp*. and its impact on epithelial cytotoxicity. **A** Microscopic images (white scale bar, 20*μm*) of germination assays on RPMI medium to automatically count and derive germination kinetics as shown in **B** for the species *A. fumigatus* (red), *A. nidulans* (blue), *A. niger* (green) and *A. terreus* (purple). **C** the relation of germination time and damage of AEC after 24h in the dynamic optimization model and parameter range with expected pathogenicity. Model prediction in **C** is compared to experimental cytotoxicity of *Aspergillus spp*. by lactatedehydrogenase (LDH) release measurements after 24h co-incubation with human A549 epithelial cells in **D** and with murine T7 epithelial cells in **E**.

Strikingly, in an experimental set-up where human (A549 cell line) and murine (T7 cell line) lung AEC are co-incubated with *Aspergillus spp*. for 24h, only those species with a fast germination show a pronounced host cell damage (see Figure 4D and E). Moreover, cytotoxicity against human cells coincides with germination speed and strongly supports model prediction. However, we observed a higher cytotoxicity of *A. fumigatus* against murine AEC while other species show comparable results (see Figure 4E). This underlines the importance of further investigations to understand differences between the human host and rodent model organisms. Further, it suggests an avoidance of elevated epithelial damage by *A. fumigatus* and its ability to hide and escape in AEC during the human immune response.

The accordance of modeling prediction with experimental data on germination as well as cytotoxicity shows that our dynamic optimization approach is able to identify key parameters of invasive aspergillosis. However, since *A. fumigatus* is the most common cause of invasive aspergillosis [10] in spite of its slightly slower germination than *A. nidulans*, the interaction of fungal pathogens with host cells and parameters defining this process represents additional decisive factors for infection outcome.

To this end, in the following part the specific roles and functions of host cells are investigated by modeling and supported by experimental investigations.

### 2.4. Neutrophils and epithelial cells primarily promote fungal clearance and cytokine release

The advantage of our model is that we can suppress a host cell population or function *in silico* to study their role and importance for infection outcome. This way we can simulate animal models with immunodeficiencies like neutropenia which is triggered by usage of cyclophosphamide and cortisone acetate in murine models of invasive aspergillosis [42]. The analysis of parameter sensitivity provides a global overview of the correlation between infection parameters and infection outcome. However, the causality of parameter influence is not explained. To this end, we performed additional *in silico* experiments with varying conidial dosages to understand and link host cell functions to infection outcome.

The conidia dose-response-curves clearly show that lack of neutrophils heavily worsens infection outcome across all dosage scenarios and host damage is barely dose-dependent (see Figures 5A and B). Further, both objectives, *i.e.*, the preservation of tissue and reduction of the fungal burden, are more impaired than by all other deficiency scenarios including lack of AM or an inhibition of cytokine release. Interestingly, lack of AM or an inhibition of cytokine release by AM show no major differences in the infection outcome compared to the reference parameter set of healthy mice (see Figures 5A and B). The optimization results for the inhibition of cytokine release by AEC or by both, AEC and AM, indicate a shift in the immune objective at low conidia dosage. In comparison to healthy mice, lung tissue is more preserved at low conidia concentrations, but with the drawback of a higher average of fungal burden during the simulated time of infection.

**Figure 5:**
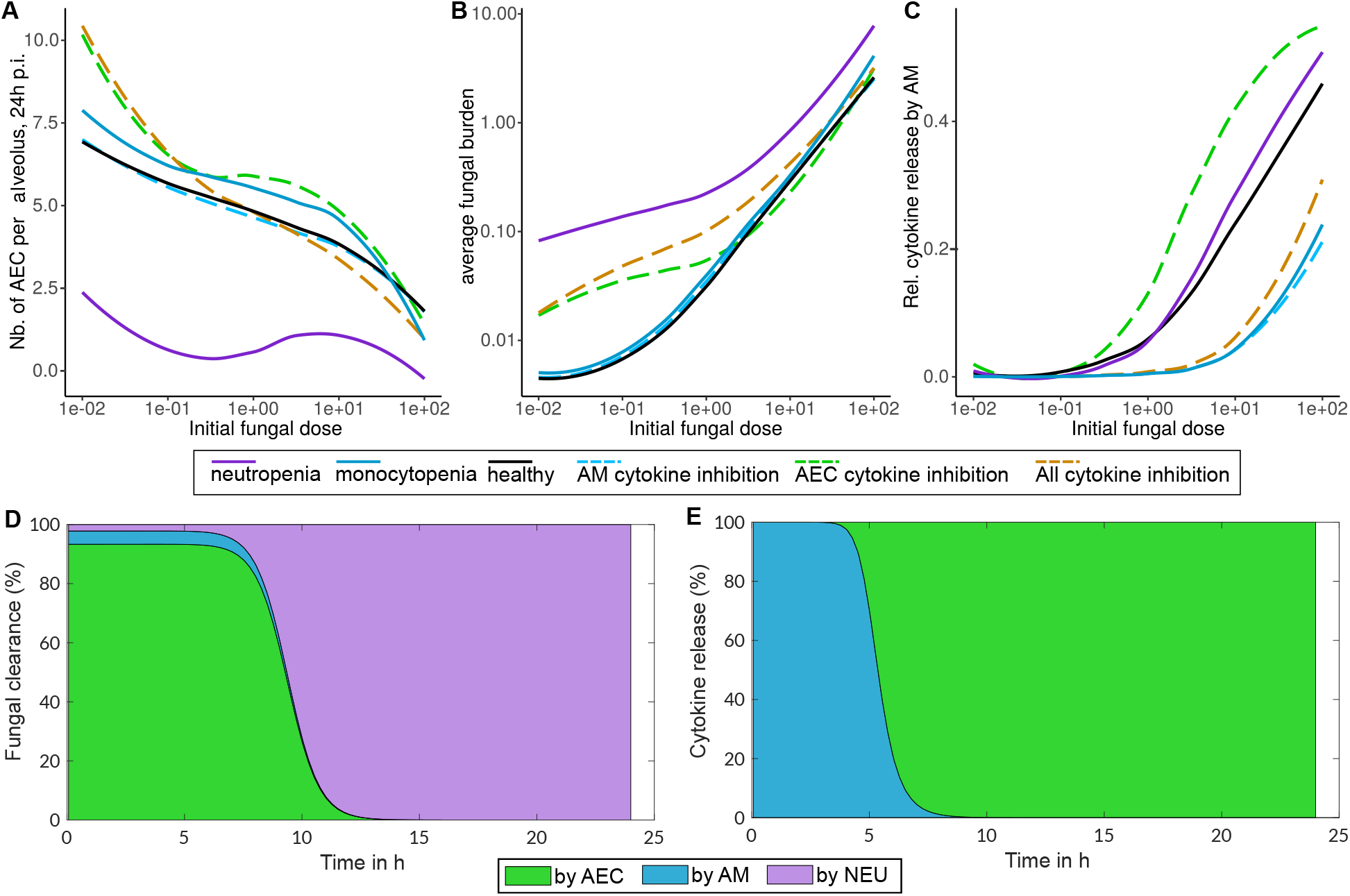
Relative contribution of host cells during infection. **A-C** Dose-response-curves with varying conidia dosages (x-axis) and the remaining number of epithelial cells at the end of simulation in **A** or the average fungal burden during simulation in **B**. The relative contribution of alveolar macrophages to cytokine release depending on the initial conidia dosage in **C**. Response curves were determined for the disturbances: lack of immune cells (1% neutrophils, solid violet; 1% macrophages solid light blue) and cytokine release inhibition (1% of cytokine release by alveolar macrophages (AM, dashed light blue), 1% release by alveolar epithelial cells (AEC, dashed green) or 1% release from both (dashed brown). **D, E** Role of macrophages (AM), neutrophils (NEU) and epithelial cells (AEC) in fungal clearance **A** and cytokine release **B** in the dynamic optimization model over time as percentage of total at each time point. Fungal clearance is calculated as the sum of killing all fungal cell types (conidia and hyphae).

In our model AM and AEC exhibit overlapping functions and duties during infection. Both are able to clear swollen conidia and to release cytokines for immune cell recruitment. However, their contribution is different depending on the initial fungal burden and during the dynamic stages of infection. When the dosage of conidia is low, the relative contribution of AM to cytokine production is nearly zero and most importantly, germination and hyphal growth are recognized by AEC (see Figure 5C). The relative contribution to cytokine release by AM rises to > 40% at very high conidia concentrations (see 5C) and indicates that AM are more important when a high number of swollen conidia is present (10-100 spores per alveolus).

This finding is supported by the relative contribution of each cell type to fungal clearance and cytokine release over the time course of infection (see Figure 5D). During infection, according to our model, AM are responsible for early recognition of conidia and AEC are the main cytokine releasing cells after germination. However, the low quantities of AM compared to AEC, lead to a much higher cytokine release by AEC compared to AM in absolute and relative terms during infection (see Figure 5E).

The prominent role of AEC in pro-inflammatory cytokine production upon fungal infection is not captured by other models of invasive aspergillosis and indicates an underestimated importance of these cells for the immune response. To quantify and support findings predicted by our model, *ex vivo* experiments with murine cells were performed and pro-inflammatory cytokines (TNF)-*α* and (IL)-6 were measured upon stimulation with *A. fumigatus* by enzyme-linked immunosorbent assay (ELISA). These cytokines were selected because of their prominent role during invasive aspergillosis in mice [41, 43–45]. Whereas AM mainly produce (TNF)-*α* after 6h of infection with conidia, murine AEC mainly produce (IL)-6 after 10h (see Figure 6A). At similar cell counts AM produce earlier (6h versus 10h), but less in absolute terms in comparison to AEC when excess release upon conidial challenge of both cytokines, (TNF)-*α* and (IL)-6, is combined (see Figure 6B). Considering the higher number of AEC than AM in an alveolus, these experimental findings strongly support our model that AEC are important mediators of the immune response.

**Figure 6:**
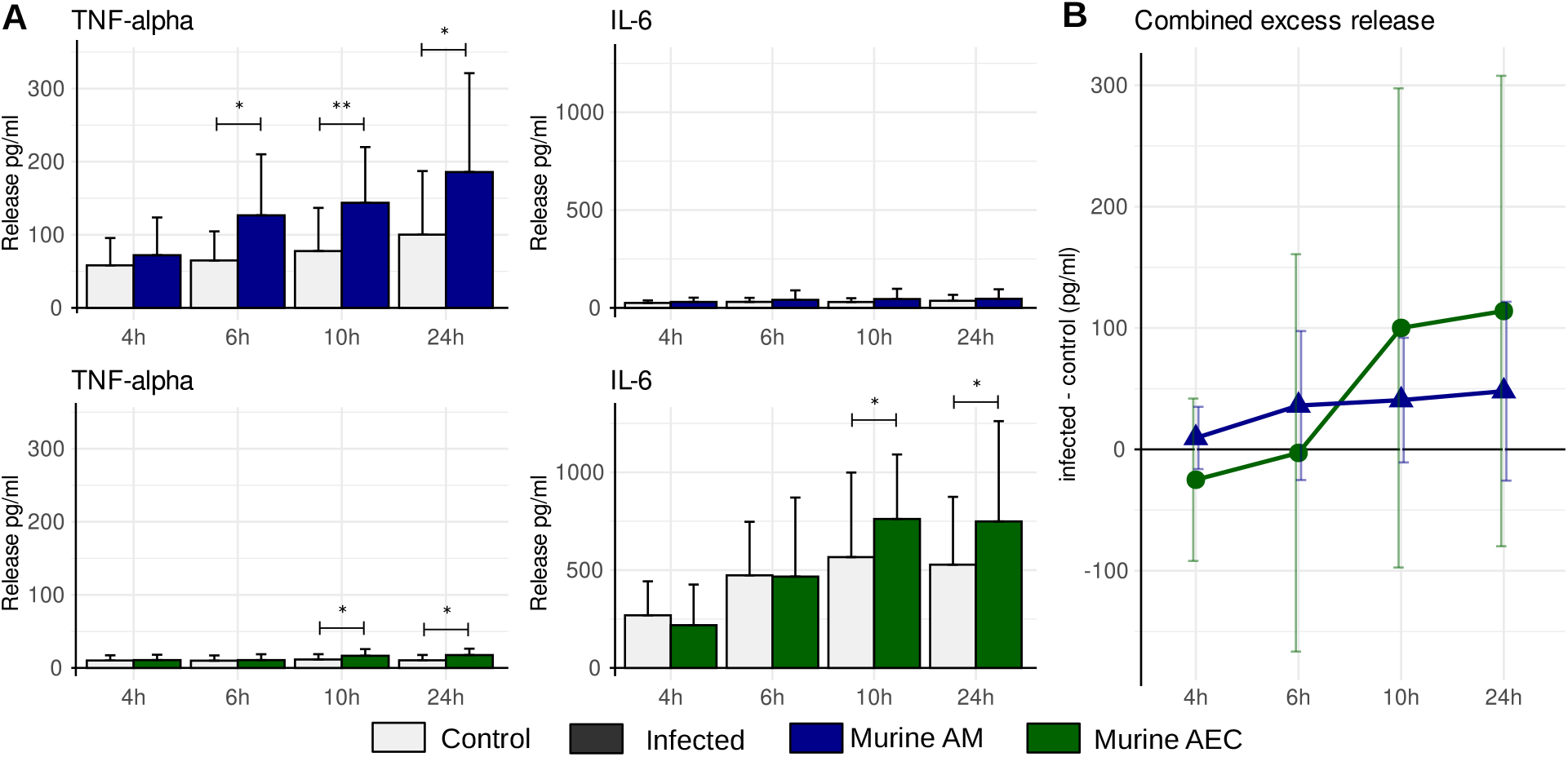
*Ex vivo* cytokine release of isolated murine alveolar macrophages (blue) and epithelial cells (green) upon infection with *A. fumigatus*. **A** Release of (TNF)-*α* and (IL)-6 over time in infected and control cells measured by ELISA. Significant differences between control and infected cells were determined by a two-tailed and paired t-test indicated by * (*p* < 0.05) and ** (p < 0.01). **B** Combined cytokine release of both (TNF)-*α* and (IL)-6 in excess over spontaneous release over time to depict contribution during early and later stages of fungal infection.

Moreover, our model reveals that AEC are important phagocytes and are predicted to be the main cell type clearing fungal spores in the first hours of infection (see Figure 5D). Nevertheless, after germination neutrophils are decisive over fungal clearance by killing hyphae. These findings highlight that alveolar epithelial fulfill a major role in fungal clearance but also in immune cell recruitment by cytokine release.

## 3. Discussion

In this study we present a unique model using the approach of dynamic optimization to study the innate immune response during invasive aspergillosis. The model’s parameterization builds on an extensive literature search and manual curation, which ensures a proper simulation of the infection dynamics. Main and distinctive features of the model are the optimization of immune cell recruitment depicting the trade-off between tissue integrity and pathogen clearance (dynamic optimization) as well as the active role of lung AEC in the immune response.

The simulations and analysis of our model revealed key parameters and insights into the roles of host cells during the innate immune response against *A. fumigatus* and other Aspergilli. We identified the state ’swollen conidia’ as most vulnerable for host defense. Hence, its time span is minimized by the fungus to escape phagocytosis and, subsequently, outcompete host cells by fast growth of hyphae. While this was indicated by experimental studies linking fungal traits with clinical observations [46, 47], we here report a quantification of this relationship. Further, we provide experimental evidence by the characterization of different *Aspergillus spp*. and their germination kinetics, as well as their cytotoxicity against human and murine AEC. Interestingly, in our and other studies *A. nidulans* shows fastest germination and a high damage to AEC. Since *A. nidulans* is not the leading cause of invasive aspergillosis in humans [48], it is likely that this species lacks immune evasion mechanisms in comparison to *A. fumigatus* [36]. However, infection by *A. nidulans* is frequently observed in chronic granulomatous disease where the lack of a fully functional NADPH-oxidases strongly limits the capability of host immune cells [49, 50]. Hence, important fungal virulence traits of *A. fumigatus* are presumably linked to an efficient escape or inhibition of phagocytosis in comparison to *A. nidulans*. This conclusion is supported by the novel observation that only *A. fumigatus* shows a stronger cytotoxicity against murine cells in comparison to human AEC (see Figure 4D and E). While this indicates an adaption of *A. fumigatus* to hide and reside in human AEC from other immune cells, further investigation is necessary to understand the differences between the human host and rodent model organisms.

As crucial host parameters for infection outcome we reaffirmed the importance of neutrophils to control filamentous growth of *A. fumigatus* which underlines that dysfunctional hyphal killing or neutropenia are major risk factors for invasive aspergillosis [28, 51]. In addition to immune effector cells, we found that lung AEC make an important part of the immune response by contributing extensively to phagocytosis of conidia and cytokine release in response to fungal germination. While this finding was partially observed and suggested in previous studies, we present here the first model which quantifies the role of AEC. Together with experimental measurements that compare the cytokine release of AM and AEC, we conclude that the role of AEC is underestimated. This conclusion is largely based on the observation that the number of AEC per alveolus is higher than the number of AM or neutrophils at early stages of infections, while phagocytosis rates and cytokine release are comparable or even higher than observed with AM (see Figure 6).

Despite their detailed investigation by experimental studies and computational models [23, 52], our results suggest that AM are involved in early recognition of swollen conidia and contribute less to fungal clearance than neutrophils and AEC. We observe a slightly higher importance of AM in low dose scenarios that indicates the importance of dosage in animal models of invasive aspergillosis to ensure transferability of results. However, our results are in line with the observation that in murine infection models neutrophil depletion, but not AM depletion, increases mortality rates [28].

Since our model focuses on the innate immune response and is a simplification of complex interactions during fungal infection, additional functions of AM are potentially not represented in the model. Such functions of AM involve *e.g.* the balance of pro- and anti-inflammatory signaling as well as the support of tissue repair [53, 54]. Further, AM link innate immunity to the adaptive immune system and are important versatile cells to maximize the robustness of the immune system [55, 56].

Additional *in silico* and *in vivo* studies are needed to fully understand the roles of AM as well as lung AEC during invasive aspergillosis. Such models and the model presented here are very valuable to identify new treatment approaches and to determine optimal treatment strategies by a combination of approaches. Dynamic optimization has been previously suggested and applied to calculate an optimal time-course of treatment protocols to combat infections by determining an optimal usage of antimicrobials and therapies boosting the immune response [57]. For such purposes, our model provides an excellent starting point to identify time-optimal treatment strategies of invasive aspergillosis.

## 4. Material and methods

### 4.1. Model formulation

We consider three different states, resting and swollen conidia (spores) as well as hyphae (filaments). Resting conidia are administered to the host’s lung at varying quantity *R*_0_. After 3-4h conidia will swell in lung alveoli and can be recognized and phagocytosed by host cells [58]. To correctly include the time needed for swelling of conidia, we model the swelling rate as a normal distribution with the mean at time point *r*_1_ = 4h and variance 1h:

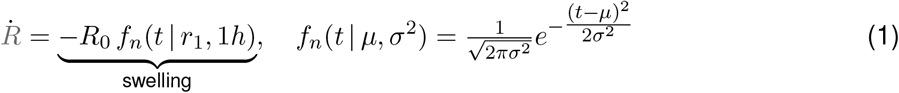

Subsequent to swelling, conidia germinate after an additional delay of *s*_1_ = 3*h* to hyphae [58, 59]. The phagocytosis by immune as well as AEC is modeled by simple bilinear terms with the specific rates (*s*_2–4_):

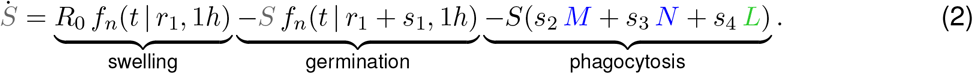

In the multicellular growth state of hyphae the number of *A. fumigatus* cells depends on the rate of germination as well as the growth rate (*h*_1_) of hyphae, which we link to the presence of AEC as a resource for growth. We model killing of hyphae by neutrophils and obtain the following description of hyphal dynamics:

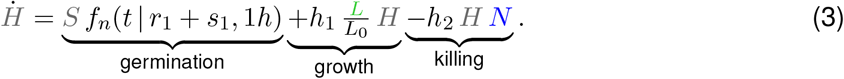

The number of AEC is influenced in our model by the lysis induced by fungal hyphae and the damage originating from active AM and neutrophils. The tissue damage by immune cells is often ignored in computational models, but is crucial to reflect recruitment and depletion of immune cells [60]. To this end, we further connect tissue damage with the pro-inflammatory cytokine level *C*:

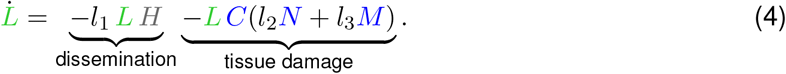

AM are of relevance for recognition and phagocytosis of swollen conidia. In addition to tissue-resident cells, more AM are recruited by transmigration and differentiation of monocytes which circulate in the blood. We assume that this recruitment has a maximal rate of *m*_1_ cells per hour and depends on the pro-inflammatory cytokine level *C* and macrophage recruitment is further optimized *via* the control variable *u*_1_(*t*). This time dependent control variable is optimized by dynamic optimization and can vary between 0 (no macrophage recruitment) to 1 (maximal recruitment) at each time point. In a similar way active depletion or deactivation of AM is modeled (*u*_2_(*t*)). Lastly, the number of AM also depends on the lysis initiated by germinating conidia and therefore we model macrophage dynamics as:

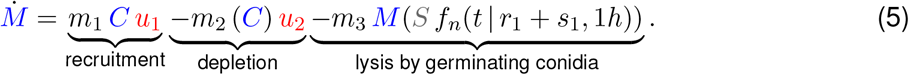

An important role of AM is the release of pro-inflammatory cytokines in response to fungal cells. However, AEC in addition release cytokines and mediate neutrophil recruitment. Thus, we model the dynamics of a pro-inflammatory cytokine level ranging from 0 (no inflammation) to 1 (maximum inflammation) as follows:

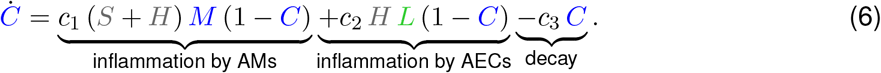

It is important to note that based on experimental observations, AEC start releasing cytokines only after conidia had germinated [30] and AM already in response to swollen conidia [7].

Although neutrophils also reside in lung tissue, they are recruited in large quantities from the blood in response to a fungal infection by pro-inflammatory cytokines like interleukin 8 [30]. The kinetics of a neutrophil population is described by:

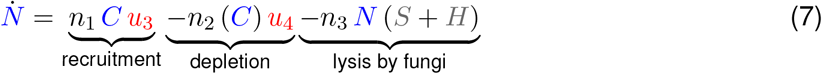

The kinetic parameters are cumbersome to be determined due to the inclusion of optimal control variables and experimental inaccessibility of *in vivo* infection parameters. Nevertheless, we carefully and extensively reviewed published experiments to estimate each parameter (see Appendix A). Parameters and cell numbers are determined per murine alveolus.

The above described ODE system and parameters are available as SBML file in Appendix B and are stored in the database BioModels [61] under the accession MODEL2105110001.

#### Constraints and objective of optimization problem

To determine the optimal innate immune response during invasive aspergillosis, the above described dynamic system has to fulfill the following constraints and objectives. Intuitively, state variables describing cell numbers are positive, pro-inflammatory cytokine level as well as control variables describing recruitment and depletion of immune cells range between 0 and 1:

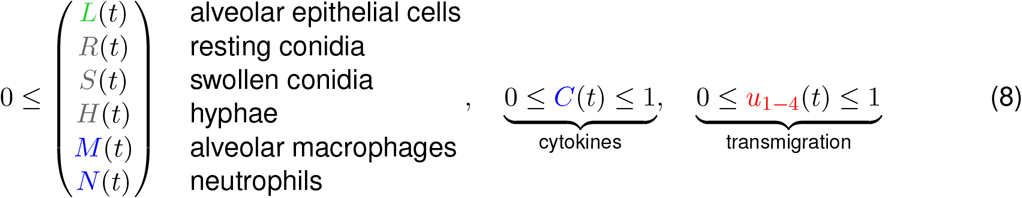

As objectives we define two main goals of the host organism. Firstly, active fungal cells (swollen conidia and hyphae) should be minimized at all time points to avert systemic infection. Secondly, unnecessary tissue damage *e.g.* by hyperinflammation and collateral damage mediated by immune cells must be avoided. We formalize as objective function:

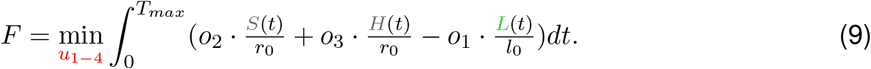

The normalization to the initial cell numbers and equal weighting *o*_1−3_ = 0.5 ensures a balanced and biological meaningful optimization result. As shown in Appendix C, considering only one of the objectives leads to undesired dynamics like continued hyphae growth or extensive tissue damage.

### 4.2. Solving the optimization problem and parameter sensitivity

The resulting dynamic optimization problem with continuous state and control variables was solved by a quasi-sequential approach established and implemented by Bartl et al. [62]. This gradient-based method has proven its capability in several previous applications to biological systems [24, 63–66] and ensures fast and robust calculation of the optimal control. To avoid local optima, we perform for each parameter set at least 100 randomizations of the initial solution, which is used to start the optimization process.

To determine the sensitivity of parameters, we sampled 500 parameter sets according to a log-normal distribution, where for each parameter the respective mode (maximum of the density function) corresponds to the literature-based parameter value (see Appendix A). The parameter sensitivity is expressed as the contribution to variance [67] and is based on the Spearman correlation *rho*(*p*) between the parameter value *p* and the objective function value:

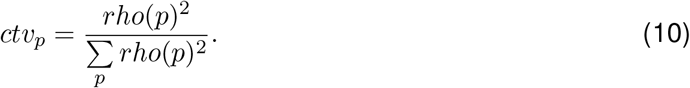

Our model enables the exploration of different scenarios like different conidia dosages and immunodeficiencies. For mice we use as initial conidia burden a dosage of 1 conidium per alveolus as ’sublethal’ and in low dose scenarios 0.1 which are comparable to *in vivo* experiments. A lack of immune cells like neutropenia or monocytopenia (lack of AM) is simulated by a reduced (1%) initial cell number and rate of recruitment. In a similar way, cytokine release inhibition is simulated by reducing the rates to 1% of the reference value.

### 4.3. Experimental evaluation of fungal and host infection parameters

#### Fungal strains and cultivation

*Aspergillus fumigatus* CEA10, *Aspergillus nidulans* FGSC A4, *Aspergillus niger* ATCC 1015 and *Aspergillus terreus* SBUG 844 were grown on *Aspergillus* minimal medium (AMM; containing 70*mM NaNO*_3_, 11.2*mM KH*_2_*PO*_4_, 7*mM KCl*, 2*mM MgSO*_4_, and 1^*μL*^/_*mL*_ trace element solution at pH 6.5) and agar plates with 1% (w/v) glucose for 5 days at 37 °C. The trace element solution was composed of 1*g FeSO*_4_ · 7*H*_2_*O*, 8.8*g ZnSO*4 · 7*H*_2_*O*, 0.4*g CuSO*_4_ · 5*H*_2_*O*, 0.15*g MnSO*_4_ · *H*_2_*O*, 0.1*gNaB*_4_*O*_7_ · 10*H*_2_*O*, 0.05*g* (*NH*_4_)_6_*M*_*o*7_*O*_24_ · 4*H*_2_*O*, and ultra-filtrated water to 1000*mL* [68]. All conidia were harvested in sterile, autoclaved water, then filtered through 30*μm* filters (MACS Milteny Biotec) and counted with a Thoma chamber.

#### Germination assay

Germination assay was performed by inoculating 1 · 10^6^ spores per *mL* in RPMI without phenol red (Thermo Fisher Scientific). At different time points pictures were taken using a Keyence BZ-X800 microscope and the number of germinated spores was determined by counting the ratio of spores undergoing germination (germ-tube formation) per field (100 cells).

#### Cytotoxicity assay

Human lung AEC A549 (ATCC-CCL 185) were maintained in F12K nutrient medium (Thermo Fisher Scientific) with the addition of 10% (v/v) fetal bovine serum (FBS) (HyClone, GE Life science) at 37°*C* with 5% (v/v) *CO*_2_. Cells of the mouse lung epithelial cell line T7 (ECACC 07021402) were maintained in F12K nutrient medium (Thermo Fisher Scientific) with the addition of 0.5% (v/v) FBS (HyClone, GE Life science) and 0.02% (v/v) Insulin-Transferin-Sodium Selenite (Sigma Aldrich). 2 · 10^5^ cells per well were seeded in 24 well plate 18*h* prior to the experiment. Before infection cells were washed once with sterile phosphate buffered saline (PBS) 1X (Gibco, Thermo Fisher Scientific) and then incubated with conidia based on the different MOIs in DMEM without phenol red (Gibco, Thermo Fisher Scientific) with the addition of 10% (v/v) FBS. A549 cells and conidia were incubated for 20*h* at 37°*C* with 5% (v/v) *CO*_2_. LDH release was measured using the CyQuant LDH cytotoxicity assay (Thermo Fisher Scientific) using the manufacturer instructions. The absorbance was determined using a Tecan Infinite 200 (LabX).

#### Isolation of alveolar epithelial type II cells and AM from mice

A total of eighteen male and female 12-18 weeks old C57BL/6J (The Jackson Laboratory) mice were used. All animals were cared for in accordance with the European animal welfare regulation and approved by the responsible federal state authority and ethics committee in accordance with the German animal welfare act (permit no. 03-072/16).

Mice were sacrificed using 125*μL* ketamine per 20*g* xylazine and the lungs were obtained as previously described [69]. After isolation, the lungs lobes were digested for 45*min* at room temperature in 1*mL* of dispase (Corning) and then the lung parenchyma was separated with the help of tweezers in 7*mL* of DMEM/F12K (Thermo Fisher Scientific) containing 0.01*mg* of DNase (Sigma Aldrich). The cell suspension was first filtered twice: through a 70*μm* and then through a 30*μm* (MACS, Miltenyi Biotec) filter and finally centrifuged. The pellet was lysed using a red blood cell (RBC) lysis buffer and re-centrifuged.

For their separation cells underwent a double magnetic labeling selection. At first, they were negatively labeled using CD45 (macrophages), CD16/32 (B/NK cells), anti-Ter (erythrocytes), CD31 (endothelial cells) (Miltyenyi Biotec) and anti-t1*α* (alveolar epithelial type I cells) (Novus Biologicals), biotin-linked antibodies. Secondly the negative fraction was collected and positively selected for alveolar epithelial type II cells using a CD326/EpCAM antibody (eBioscience). Both labeling steps were perfomed at 4°*C* for 30*min*, and followed by a second labeling with Anti-Biotin Microbeads Ultrapure (Miltenyi Biotec) for 15*min* at 4°C. The separation was performed using an autoMACS Pro Separator machine (Miltyenyi Biotec). The final cell suspension containing type II AEC was resuspended in mouse tracheal epithelial cells (MTEC) basic medium [70]: DMEM-F12K +1% HEPES, Na-bicarbonate, L-glutamine, penicillin-streptomycin, 0.1% amphotericin B (Gibco) supplemented with 0.001% Insulin-transferrin (Gibco), 0.1*^μg^/_mL_* cholera toxin (Sigma Aldrich), 25*^ng^/_mL_* epidermal growth factor (Invitrogen) and mice fibroblast growth factor 7 (R&D system), 30*^μg^/_mL_* bovine pituitary extract (Gibco), 30*^ng^/_mL_* multilinear hemopoietic growth factor, 50*^ng^/_mL_* human fibroblastic growth factor 10 (R&D system), 5% FBS, 0.01*μM* retinoic acid (Sigma Aldrich) and 10*μM* Rho kinase inhibitor (ROCK) (Thermo Fisher Scientific).

The isolation of AM was performed similarly, only using the first labeling step with CD45 antibodies. AM were re-suspended in MTEC basic medium with the addition of 10% (v/v) FBS.

From each mouse, a total of 1 · 10^6^ cells were seeded onto 8 well Millicells slides (Merc, Millipore), pre-coated with 100*^μg^/_mL_* of fibronectin (Sigma Aldrich), for AEC, and incubated at 37°C and 5% (v/v) *CO*_2_. The medium was changed every 2 days and the cells were left to rest for 7 days prior the experiment.

#### Infection with A. fumigatus and cytokine measurement

Alveolar epithelial type II cells were infected with *A. fumigatus* CEA10 conidia at a multiplicity of infection (MOI) of 5, for 4, 6, 10, and 24h. At these time points the supernatant was collected, centrifuged at 300 · *g* for 5 min and then stored at −20°C for 24h until measurements. The levels of interleukin (IL)-6 and tumor necrosis factor (TNF)-*α* were detected using ELISA kits (Biolegend) following the manufacturer’s instructions.

## Acknowledgments

We acknowledge support by the Deutsche Forschungsgemeinschaft (DFG) CRC/Transregio 124 ‘Pathogenic fungi and their human host: Networks of interaction’ (support code 210879364) sub project B1 (J.E. and S.S.) and A1 (F.R. and A.B.). Further, C.K. acknowledges support by the Cluster of Excellence ‘Precision Medicine in Chronic Inflammation’ (support code EXC 2167) and L.R. by the Cluster of Excellence ‘Balance of the Microverse’ (support code EXC 2051 - Project-ID 390713860). We thank Elina Wiechens for an initial parameter search in the literature and thank Pedro Moura-Alves, MPIIB Berlin, for providing us the mouse T7 alveolar lung cells. In addition, we thank Dr. Maria Straßburger for the help with mouse experiments.

## Appendix A. Parameter estimation

Documentation of parameter estimation and calculation

*L*(0),*L*_0_ Average number of AEC (type I and II) per alveolus. The total number of AEC in several mammalian species was elucidated by Stone et al. [38] and reports for murine lungs 11.6 + 14.8 = 26.4 · 10^6^ *cells* (type I and II). Knust et al. [39] determined 2.31 · 10^6^ alveoli for mice with a comparable weight (20 *g*) and we estimate 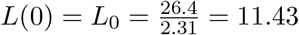 *cells*.
*l*_1_ Rate of hyphae destroying AEC. In [71], epithelial damage was assessed in mice by LDH release in bronchial lavage showing a 2.5 fold change of damage after 24h. The rate is calculated by 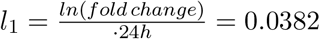.
*l*_2_ Rate of neutrophils damaging AEC. In [72], neutrophils were stimulated *in vivo* with LPS and cell damage in lungs was assessed as free elastase activity in BALF. The relative damage was 72% after 12h and we estimate 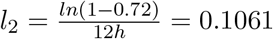
*l*_3_ As *l*_2_ but for AM. Host damage mediated by AM is less frequently measured and we therefore rely on data of Hirano [73]. In that study rat epithelial cells (SV40T2) were co-cultured with rat alveolar macrophages and stimulated by LPS. The damage was determined by transepithelial resistance and dropped from 11 Ω*cm*^2^ to 0.5 Ω*cm*^2^ within 24h. To this end, we calculate 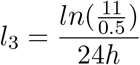.
*r*(0), *R*_0_ Initial dose of resting conidia. Value depends on the scenarios. *In vivo* experiments with mice typically challenged with 10^5^-10^7^ conidia. Therefore, alveolus occupation is typically 0.1 - 10 conidia per alveolus. A detailed calculation is also provided by Blickensdorf et al. [23].
*r*_1_ Average time for swelling of conidia. After 3h the conidial rodlet layer is disrupted [58].
*s*_1_ Average time for germination of swollen conidia. Total time from resting conidia to germination is on average 7h [59]. Therefore *s*_1_ = 7*h* – *r*_1_ = 4*h*.
*s*_2_ Killing rate of swollen conidia by AM. Intensively studied, but rates show high variation across studies [26]. In contrast, the killing rate of conidia by AEC is more rarely calculated. However, in [29] both was investigated. Accordingly, murine AM showed faster killing than AEC along all timepoints. For murine AM, conidia survival was 0.6% after 12h of interaction and we calculate 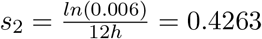.
*s*_3_ Killing rate of swollen conidia by neutrophils. Phagocytosis and killing rate of conidia by neutrophils have been frequently studied. However, in the study of Alflen et al. [74] the killing rate of conidia and hyphae was compared and murine neutrophils were used. Conidial viability dropped to 39% after 4h of co-incubation and we calculate 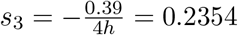.
*s*_4_ Killing rate of swollen conidia by AEC. From the same study as for s_2_ [29], AEC possessed a slower killing rate and we calculate 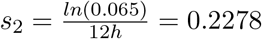.
*h*_1_ Growth rate of hyphae. The increase of hyphal length with time is a good estimator for doubling times of cells since cell size is rather constant. In glucose medium and at 37°C the length of hyphae doubles every 1h [75]. The rate is then 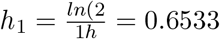.
*h*_2_ Killing rate of hyphae by neutrophils. From the same study as used to estimate *s*_3_ [74], the viability of hyphae was measured after co-incubation with neutrophils. After 4h 68% of initial fungal cells were viable, which equals a rate of 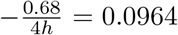. However, hyphae are growing at the same time and therefore we estimate *h*_2_ = *h*_1_ +0.0964 since neutrophil killing outpaces hyphal growth by the above calculated rate.
*M*(0) Number of resident AM per alveolus in an uninfected host. Recently, Amich et al. [40] were able to use 3D light sheet fluorescence microscopy to investigate invasive aspergillosis in *in vivo* mouse models. Due to this imaging approach one can precisely determine the number of immune cells within an alveolus. On average they find *M*(0) = 0.2926 AM per alveolus. Note, that in the agent-based model of Blickensdorf et al. [23] a slightly higher number of AM per murine alveolus (0.74) was calculated based on whole tissue cell counts [38].
*m*_1_ Rate of recruitment of AM by influx and maturation of monocytes. To complement data used for *M*(0) and *N*(0). we estimate the recruitment rates of AM and neutrophils based on data from a previous publication [41] of the same group of authors, which performed 3D light sheet fluorescence microscopy [40]. In that publication a similar murine infection model of invasive aspergillosis was used, but cell numbers were quantified by fluorescence-activated cell sorting (FACS) of lung tissue samples. For AM a rise from 2.75 to 910^5^*cells* in 4h after a mild dose of conidia was reported. We estimate 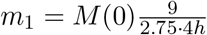.
*m*_2_ Rate of alveolar macrophage depletion. Based on the same study as *m*_1_ [41] macrophage level was on control level after 12h of peak inflammation. So we calculate 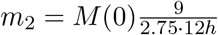.
*m*_3_ Lysis rate of AM by swollen conidia. While phagocytosis and killing rate by AM are frequently reported the *vice versa* scenario is rarely depicted. For mice cytotoxicity of swollen conidia was reported [76] to be 84% after 24 h that corresponds to a rate of 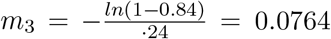.
*c*_1_ Release rate of pro-inflammatory cytokines by AM. We intentionally do not model a single cytokine due to the complex network of cytokine functions and signaling to recruit neutrophils. Based on several studies investigating the time course of prominent pro-inflammatory cytokines we decided to set the cytokine release rate to *c*_1_ = 1 to mimic the peak in cytokine concentration at 5-10h after infection or stimulation with LPS.
*c*_2_ Release rate of pro-inflammatory cytokines by lung AEC. Cytokine release of lung AEC is not well understood and only covered by few studies. A pioneering study [77] investigated the potential of human lung AM and AEC to recruit neutrophils when stimulated by LPS. Since AEC showed a 43.75% higher number of recruited neutrophils under similar conditions, we set *c*_2_ = 1.14375.
*c*_3_ Decay rate of pro-inflammatory cytokines. As for *c*_1_ we estimated the decay rate based on the investigation of several time courses and observed that decay is slower than release. For reference we set *c*_3_ = 0.1.
*N*(0) Number of tissue resident neutrophils. As described for *M*(0) microscopy data of murine lungs reported a neutrophil density of *N*(0) = 0.2628 neutrophils per alveolus [40].
*n*_1_ Rate of neutrophil recruitment by influx of leukocytes. Similar to the definition of *m*_1_ an increase of neutrophils is reported in [41] from 1.9 · 10^5^*cells* to 7.6 · 10^5^cells. To this end, we calculate 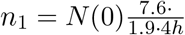.
*n*_2_ Rate of neutrophil depletion. Based on the same study as *n*_1_ [41] neutrophil level was on control level after 12h of peak inflammation. So we calculate 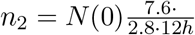.
*n*_3_ Lysis rate of neutrophils by fungal cells. In [78] induced decondensation of nuclei of neutrophils (1 · 10^6^*cells*) was measured in response to short hyphae (750*CFU* with length of 10 - 100*μm*). After 6h of co-incubation 28 % of neutrophil nuclei were decondensed. Based on the assumption that hyphae contain around 10 nuclei at the reported length, we estimate an MOI of 1:10 and calculate the rate as 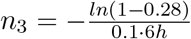.

## Appendix B. SBML model description

The described model of invasive aspergillosis is provided as SBML and COPASI file. To enable simulation, control variables *u*_1–4_ are fixed. Further, the model is archived in EBI BioModels under the accession MODEL2105110001.

## Appendix C. Influence of weighting innate immune response objectives

Model behavior with no fungal burden minimization and only minimization of tissue damage.

**Figure C.7:**
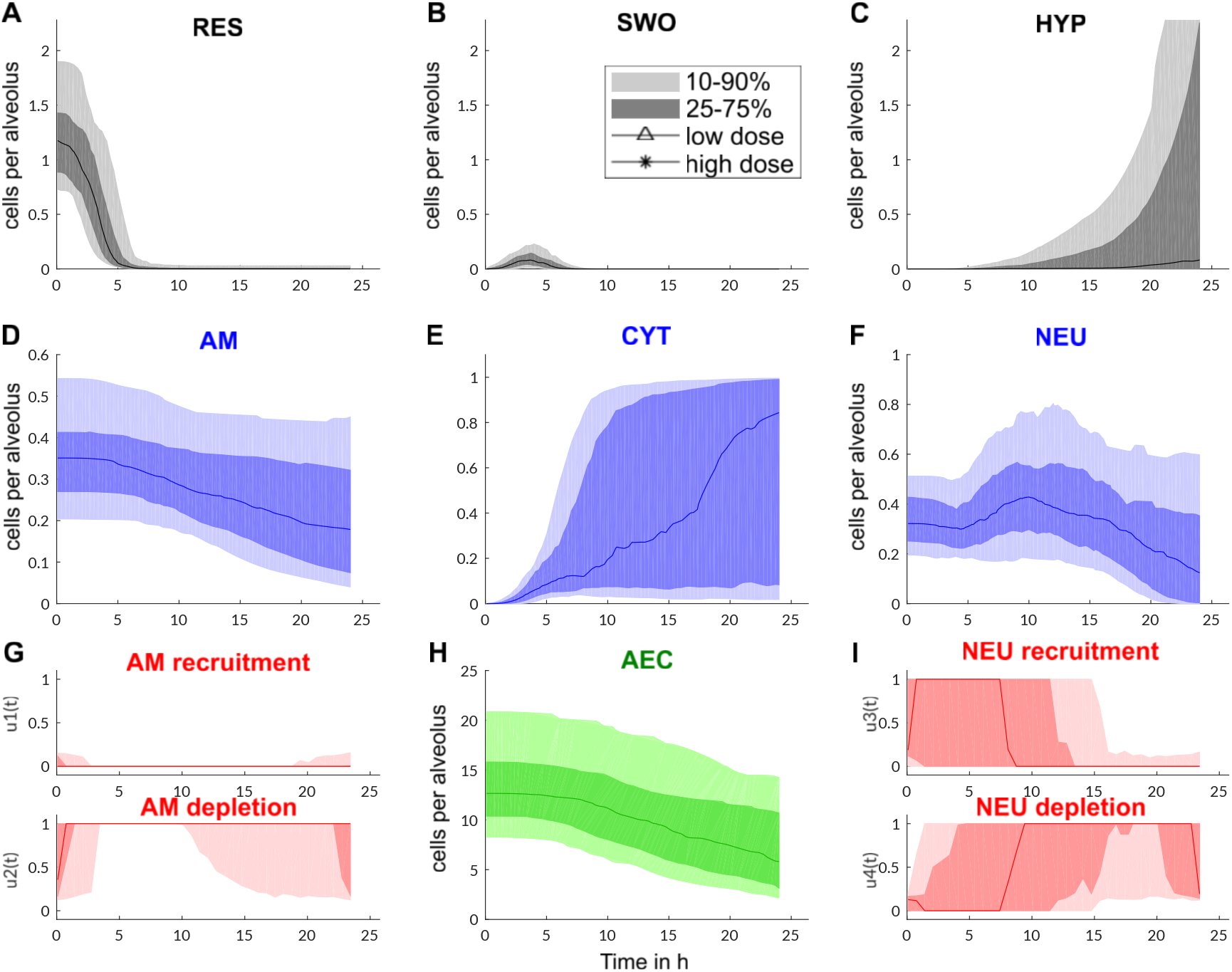
Dynamics of innate immune response of the murine host challenged with a high dose of conidia (one per alveolus). The solution space of 100 parameter sets is depicted with shadings indicating the confidence intervals of time courses. Colors as in previous figures and equations.

Model behavior with only fungal burden minimization and no minimization of tissue damage.

**Figure C.8:**
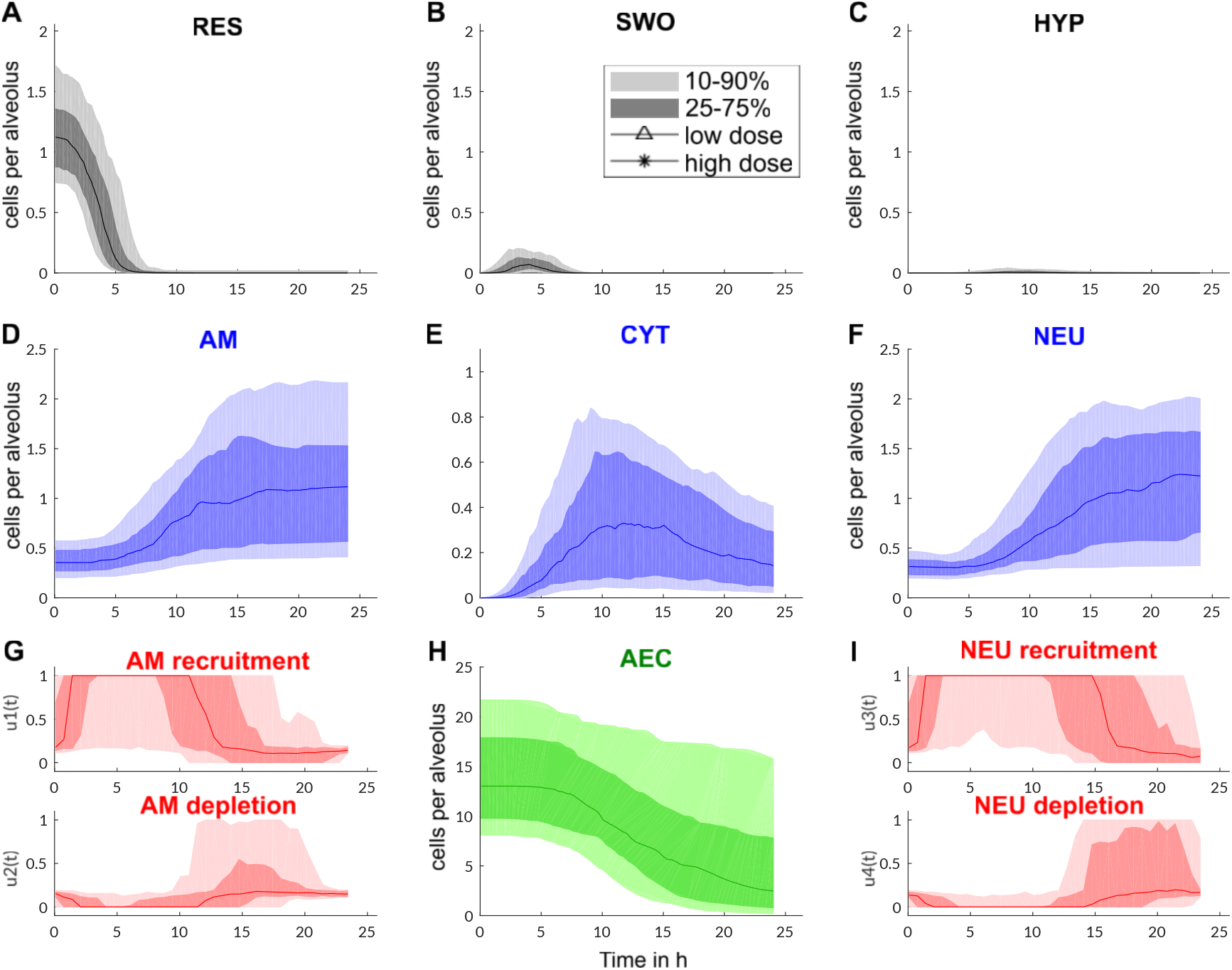
Dynamics of innate immune response of the murine host challenged with a high dose of conidia (one per alveolus). The solution space of 100 parameter sets is depicted with shadings indicating the confidence intervals of time courses. Colors as in previous figures and equations.

**Figure C.9:**
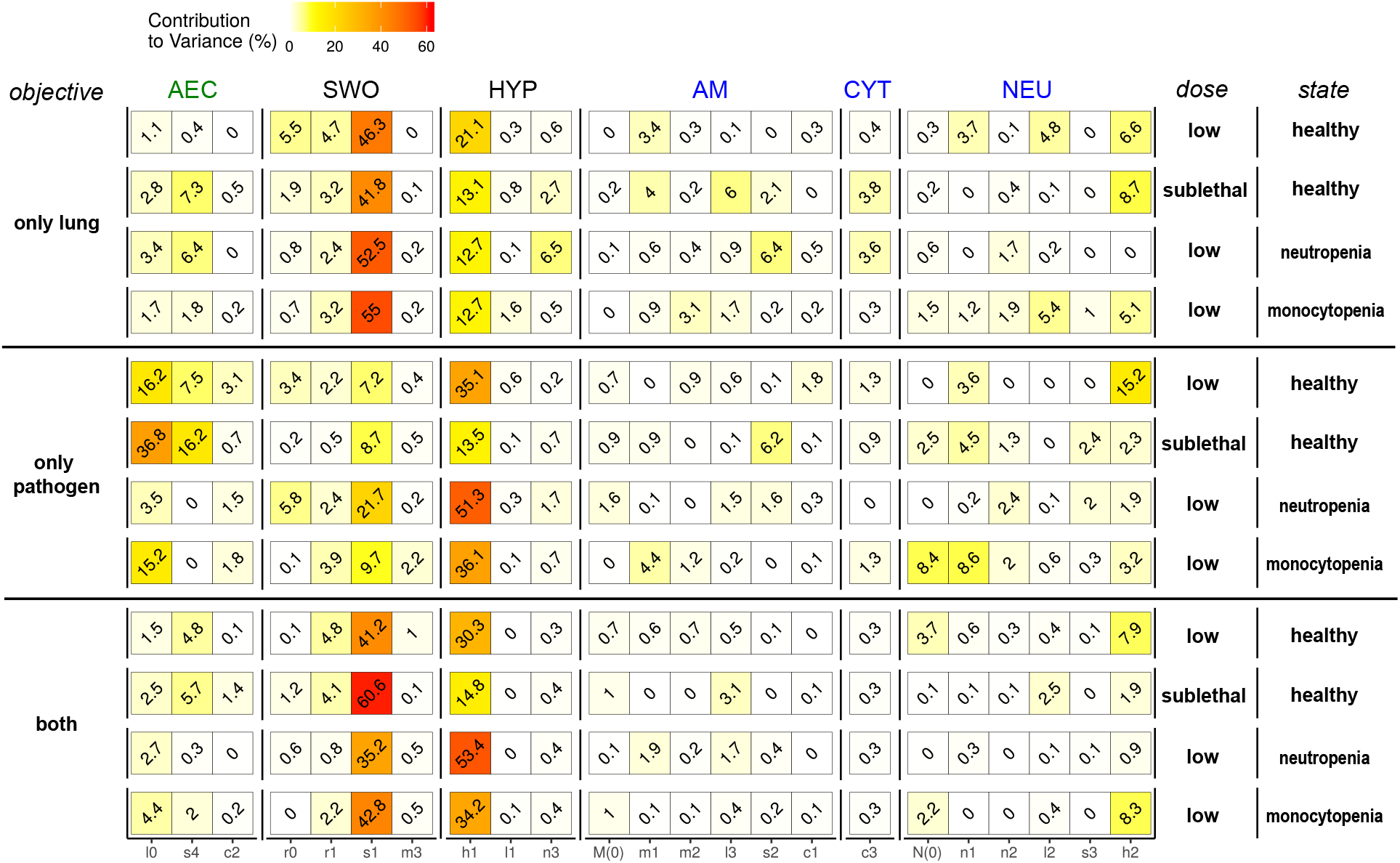
Influence of parameters on the severeness of infection depicted by the contribution to variance (brown). This relative contribution is based on Spearman rank correlation of parameter value and objective value of the optimal solution.

## Appendix D. Influence of MOI on cytotoxicity of *Aspergillus spp*. against epithelial cells

**Figure D.10:**
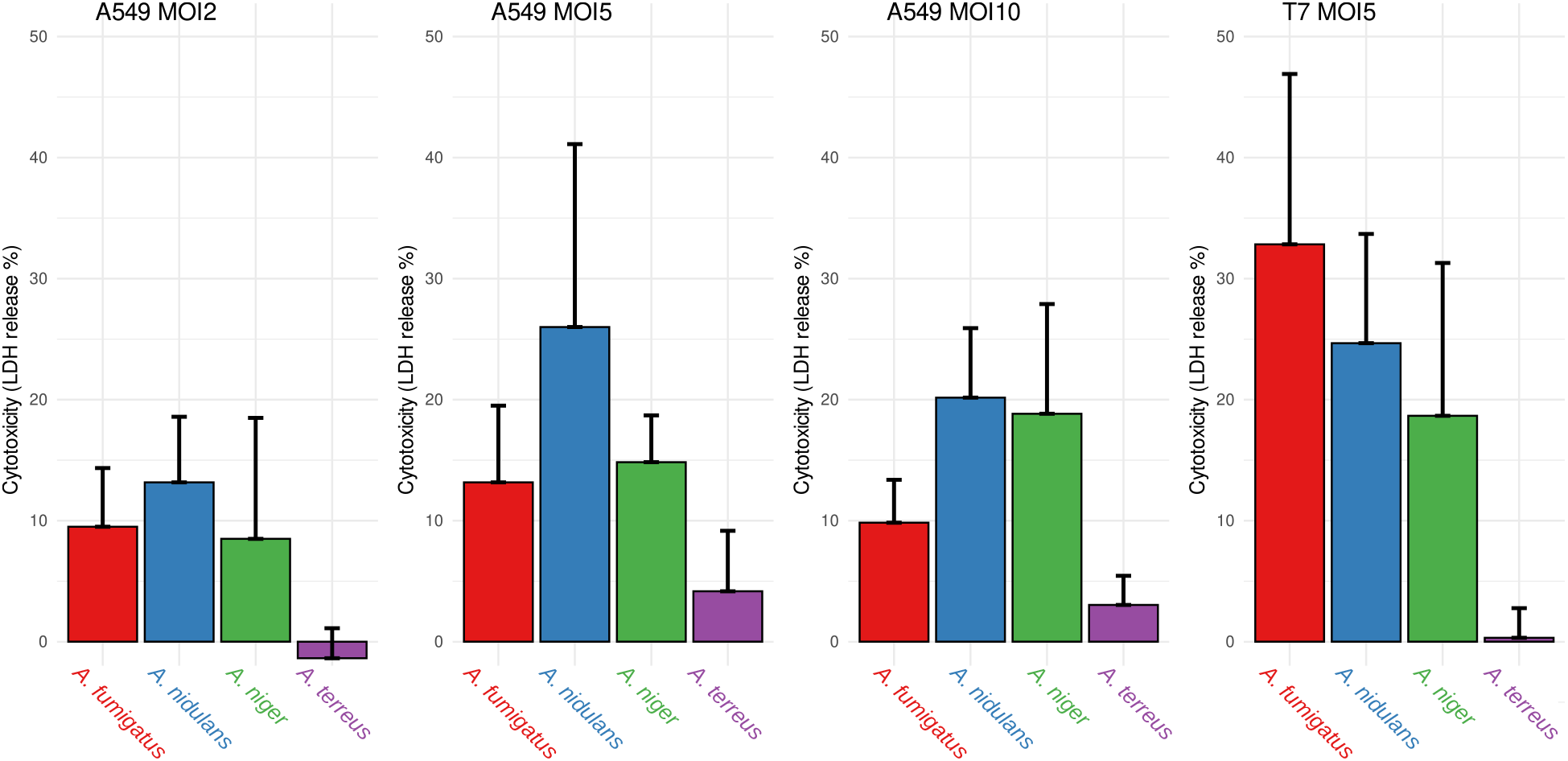
In addition to the depicted LDH release measurements of epithelial cells upon 24h co-incubation with *Aspergillus spp*. at an *MOI* = 5 (main text Figure 4D and E), cytotoxicity was measured for *MOI* = 2 and *MOI* = 10 for human A549 cells to demonstrate the influence of fungal burden.

## Notes

### Competing Interest Statement

The authors have declared no competing interest.

